# CCG interruptions are unstable and hypermethylated in DM1 patients

**DOI:** 10.1101/2024.10.10.617614

**Authors:** Sonia Lameiras, Eirini Maria Lampraki, Duncan Kilburn, Tina Alaeitabar, Ismail Jamail, Nicolas Servant, Sylvain Baulande, Sarah Kingan, Sam Holt, Valeriya Gaysinskaya, Guilherme De Sena Brandine, David Stucki, Tanya Stojkovic, Guillaume Bassez, Denis Furling, Geneviève Gourdon, Stéphanie Tomé

## Abstract

Myotonic dystrophy type 1 (DM1) exhibits highly heterogeneous clinical manifestations caused by an unstable CTG repeat expansion reaching up to 4,000 CTG. The dynamics of CTG repeats and clinical variability depends on the CTG repeat number, CNG repeat interruptions, DNA methylation, somatic mosaicism, but also gene modifiers. Around 10% of the DM1 population carries triplet repeat interruptions (CCG, CGG, CTC, CAG), which differ in number and nature between DM1 families. CCG interruptions have been associated with stabilization of the CTG repeat expansions and a milder phenotype in the DM1 population. However, the dynamics and precise role of interruptions in CTG repeat instability remain relatively underexplored due to the complexities of analyzing them. In this study, we showed that the number of CCG interruptions within the expansion varies within tissues and between generations using single-molecule real-time long-read sequencing. Although the interrupted expanded alleles showed global stabilization, the CCG interruptions themselves displayed variability across generations and within somatic tissues. Importantly, our findings reveal, for the first time, CCG hypermethylation within the CTG expansion, which is linked to downstream hypermethylation of the repeat. These results support the hypothesis that methylation of CCG interruptions within the expanded *DMPK* alleles may contribute to the stabilization of trinucleotide repeats.

## Introduction

Short tandem DNA repeats represents an important source of genetic variation and phenotypic diversity in both health and disease contexts. However, their complex nature poses significant challenges to fully understanding and characterizing their implication in human genetic disorders. Among these repeat elements, unstable repeat expansions are associated with more than sixty diseases, including myotonic dystrophy type 1 (DM1) (1). DM1 is the most complex and variable disease caused by an unstable CTG repeat expansion in the dystrophia myotonica protein kinase (*DMPK*) gene, with highly heterogeneous clinical manifestations. Larger expansions are associated with more severe symptoms and a decrease in age of disease onset (2). DM1 family pedigrees have shown that over 90% of transmissions lead to expansions of the CTG repeats, while approximately 10% result in contractions (3).

Various model systems have significantly advanced our comprehension of the molecular mechanisms involved in the formation of trinucleotide repeat expansions. Trinucleotide repeat instability depends strongly on the length and nucleotide sequence of the repeat expansions, as well as by the secondary structures formed by the triplet repeats (4). Unconventional DNA structures, including hairpins and RNA-DNA hybrids (R-loops), which arise during *DMPK* transcription, serve as mutagenic intermediates in CTG repeat expansions (5). The intricate mechanisms involved encompass DNA replication and repair machinery, as well as epigenetic processes. CAG/CTG repeats decelerate replication forks in a length-dependent manner, leading to aberrant DNA replication that destabilizes triplet repeats in DM1 fibroblasts and *in vitro* models (6–8). Furthermore, MSH2 and MSH3, two mismatch repair proteins essential for maintaining genomic integrity, play a pivotal role in triplet repeat expansions *in vivo* (9–15). In DM1 human fibroblasts, the instability of CTG repeats tends to expand following genome-wide demethylation induced by DNA methyltransferase inhibition (16). This suggests that DNA methylation may modulate CTG repeat instability in DM1 patients by either inducing methylation changes at the DM1 locus or elsewhere in the genome, thereby impacting the expression of gene modifiers associated with CTG repeat instability. Recent finding suggests that DNA methylation levels were associated with variability in somatic mosaicism in DM1 patients, based on a pyrosequencing approach (17). Although numerous studies have provided insight into the mechanisms of CTG repeat expansions, the mechanisms that generate contractions or stabilize trinucleotide repeats remain less well-described. The majority of DM1 patients inherit a pure CTG repeat expansion. However, up to 10% of the DM1 population carries triplet repeat interruptions (CCG, CGG, CTC, CAG), which differ in number and/or type between DM1 patients (18). These interruptions are associated with stabilization and/or contractions of the CTG repeat expansion, often leading to milder symptoms in the DM1 population. This suggests that sequence interruptions are an important modifier of the CTG repeat instability and DM1 phenotype (19–28). Various studies have demonstrated that interruptions such as AGG, CAT, GGA, and CCG, as described in spinocerebellar ataxia type 1, fragile X syndrome, Friedreich ataxia and DM1, can affect DNA secondary structure formation, nucleosome positioning, methylation status, and transcript levels (29–34). The aberrant formation and/or stability of secondary structures by sequence interruptions can impact CTG repeat instability by altering DNA repair and/or replication fork progression. Additionally, destabilization of interrupted CTG repeats has been observed in mismatch repair-deficient yeast, suggesting that mismatch repair proteins such as MSH2 are crucial for stabilizing the interrupted CTG sequence in this model (35). Despite these findings, the role of interruptions in triplet repeat stabilization/contraction, as well as their consequences on pathogenesis and clinical outcomes, remains poorly understood due to limitations in the standard procedures for analyzing them.

Third-generation long-read sequencing technologies offer an unprecedented opportunity to decipher large, complex regions of the human genome and reveal novel structural variants (36). Previously, our lab has shown that the Single-Molecule Real-Time (SMRT) long-read sequencing (LRS) (Pacific Biosciences) is a powerful tool for measuring CTG repeat size, identifying structural variants and estimating somatic mosaicism in DM1 (37). To complete our previous study, we used free amplification long-read sequencing in DM1 atypical patients carrying a CCG-interrupted expanded allele to precisely analyze the behavior of CCG repeats in blood and other tissues, as well as the methylation status of the expansions. In this study, the single-molecule approach reveals that the number of CCG interruptions within the expansion varies across tissues and between generations, despite the globally stable behavior of the interrupted expanded alleles. Moreover, we have demonstrated at the molecular level that CCG interruptions within the expansion are methylated and are associated with hypermethylation of the flanking repeat sequence. These findings support the hypothesis that methylation of CCG interruptions within the *DMPK* expanded alleles could induce stabilization of the trinucleotide repeats.

## Materials and Methods

### DNA samples from DM1 patients

DM1 patients were recruited by the Neuromuscular Disease Reference Center at Pitié-Salpêtrière Hospital in France and the DM-Scope registry (38). Written informed consent was obtained from all participants. Blood samples were collected in EDTA tubes at the Neuromuscular Disease Reference Center. Genomic DNA was extracted using either the Chemagic™ 360 instrument, which utilizes PerkinElmer’s patented magnetic bead technology, or the Nanobind kit from Pacific Biosciences (https://www.pacb.com/wp-content/uploads/Procedure-checklist-Extracting-HMW-DNA-from-human-whole-blood-using-Nanobind-kits.pdf). Lymphoblastoid cell lines and fibroblasts were provided by the DNA and cell bank at Genethon and Anger Hospital, respectively.

### Generation of CTG repeats amplicons

The normal and expanded CTG repeat alleles were amplified by PCR using either 0.4 μM ST300F (5ʹGAACTGTCTTCGACTCCGGG-3ʹ) and ST300R (5ʹ-GCACTTTGCGAACCAACGAT-3ʹ) primers or barcoded ST-300 primers, as per the protocol outlined by Mangin et *al.* in 2021 (39). Briefly, 5 ng of DNA was amplified in a 25-μL reaction using 0.4 μM primers, 1× Custom master mix (Thermo Fisher Scientific, Courtaboeuf, France) and 0.04U ThermoPerfect 250u (Genelab, UK; GL-801). The following cycling conditions were used: 5 min at 96 ◦C; 45 s at 96 ◦C, 30 s at 60 ◦C and 3 min at 72 ◦C (30 cycles); 1 min at 60 ◦C and 10 min at 72 ◦C (1 cycle). After amplification, PCR products for each sample were pooled and purified using the 0.5X AMPure PB beads (Pacific Biosciences, Menlo Park, CA, USA) clean-up procedure. AMPure PB beads were employed to eliminate unbound primers and PCR products corresponding to the *DMPK* normal allele.

### Library preparations and long-read sequencing

#### Amplicon approach

SMRTbell libraries were prepared from 2μg of pooled amplicons, using the SMRTbell Prep Kit 3.0, following PacBio’s Procedure & Checklist « Preparing multiplexed amplicon libraries using SMRTbell® prep kit 3 ». Briefly, this protocol includes a first step of end-repair and a-tailing to enable the following ligation of PacBio adaptors at each amplicon extremities. Then a nuclease treatment was applied to digest incomplete SMRTbells. After purification with SMRTbell Cleanup beads, final libraries quantification and quality assessment was performed using Qubit fluorometric assay (Invitrogen) with dsDNA High Sensitivity Assay Kit and Bioanalyzer 12000 kit (Agilent). Binding was performed with the Sequel II Binding Kit 3.1. Sequel II System run conditions included a 2h immobilization time, a 0.8h pre-extension and 30h movie time per SMRT Cell.

#### Amplification-free targeted approach

SMRTbell libraries were prepared from 2μg of genomic DNA, using the SMRTbell Prep Kit 3.0, following PacBio’s Procedure & Checklist « Generating PureTarget™ repeat expansion panel libraries » (https://www.pacb.com/wp-content/uploads/Application-note-Comprehensive-genotyping-with-the-PureTarget-repeat-expansion-panel-and-HiFi-sequencing.pdf) (40). This protocol consists in a first step of gDNA dephosphorylation, to remove off-target molecules in the final product. The gRNA/Cas9 complex was then prepared and dephosphorylated genomic DNAs were digested at the targeted regions. After A-tailing step, barcoded adapters were ligated to the ends of each targeted DNA fragment. Failed ligation products and gDNA fragments were removed by nuclease treatment. Libraries were then pooled (with equal volume) to be purified with SMRTbell Clean up beads. An additional step of wash was applied with SMRT boost beads to prepare the libraries for sequencing. Binding was performed with the Sequel II Binding Kit 3.2 following PacBio’s specific recommendations for PureTarget libraries. Sequel II System runs conditions included a 4h immobilization time, no pre-extension, 30h movie time per SMRT Cell and kinetic information to be retained.

Bioinformatics analysis

HiFi reads sequenced on a Sequel IIe which performed CCS analysis onboard using v6.3 of the CCS tool. The instrument also carried out 5mC calling at all CpG sites using primrose v1.9.0 with default parameters. HiFi reads (those attaining quality of ≥Q20) were mapped to the human genome reference version human_GRCh38_no_alt_analysis_set using an automated pipeline in PacBio SMRT Link software (v11.1), based on pbmm2 v1.9.0 in -CCS mode. Identification of reads which spanned the *DMPK* locus was performed using TRGT software v0.3.4 with default settings. Spanning reads were visualised using TRVZ software, which offers modes for visualising loci based on sequence motifs or based on methylation status. Reads were demuxed with the SYMMETRIC-ADAPTERS preset using lima v2.9.0. Demuxed reads were mapped to hg38 reference genome using pbmm2 v1.13.1 using the HIFI preset. Alleles were genotyped and reads were assigned to each allele using TRGT v0.9.0 using the flags: -- genotyper=cluster --aln-scoring=1,1,0,1,1,1000 --min-flank-id-frac=0.7 --flank-len=200 -- output-flank-len=200 --max-depth=10000 --min-quality=-1.0 (41).

## Results

### 1- CCG repeat number varies between generations and within tissues

In this study, a new DM1 family was recruited based on clinical presentations, specifically targeting individuals exhibiting atypical symptoms without reported severe cardiac, respiratory, or muscular abnormalities. These clinical features deviate from the typical presentation observed in DM1 patients of similar age and expanded allele size (42). First, we characterized the DM1 locus in this family by performing long-read amplicon sequencing on blood samples from individuals #17538, #15841, and #17623 (Table 1 and Figure 1). In these patients, we identified a series of CCG interruptions in the middle of the expansion, representing approximately 13% of the expansion in the blood of individuals #17538 and #15841, and 29% in individual #17623 (Table 1 and Figure 1). The estimated sizes of expansion in #17538, #15841, and #17623 are 405, 369, and 276 repeats, respectively (Figure 1). In this family, the CCG-interrupted expanded allele is associated with the absence of an increase in mutated allele size across generations and no worsening of symptom severity. Interestingly, single-molecule amplicon sequencing reveals that the number of CCG interruptions varies among the amplified DNA molecules in the blood samples from individuals #17538, #15841, and #17623, ranging from 0 to 241, 215 and 236, respectively. The percentage of CCG interruptions increases between individual #15841 and #17623, suggesting a rising trend of CCG interruptions across male transmission despite a stabilization of expansion in this family.

**Figure 1:**
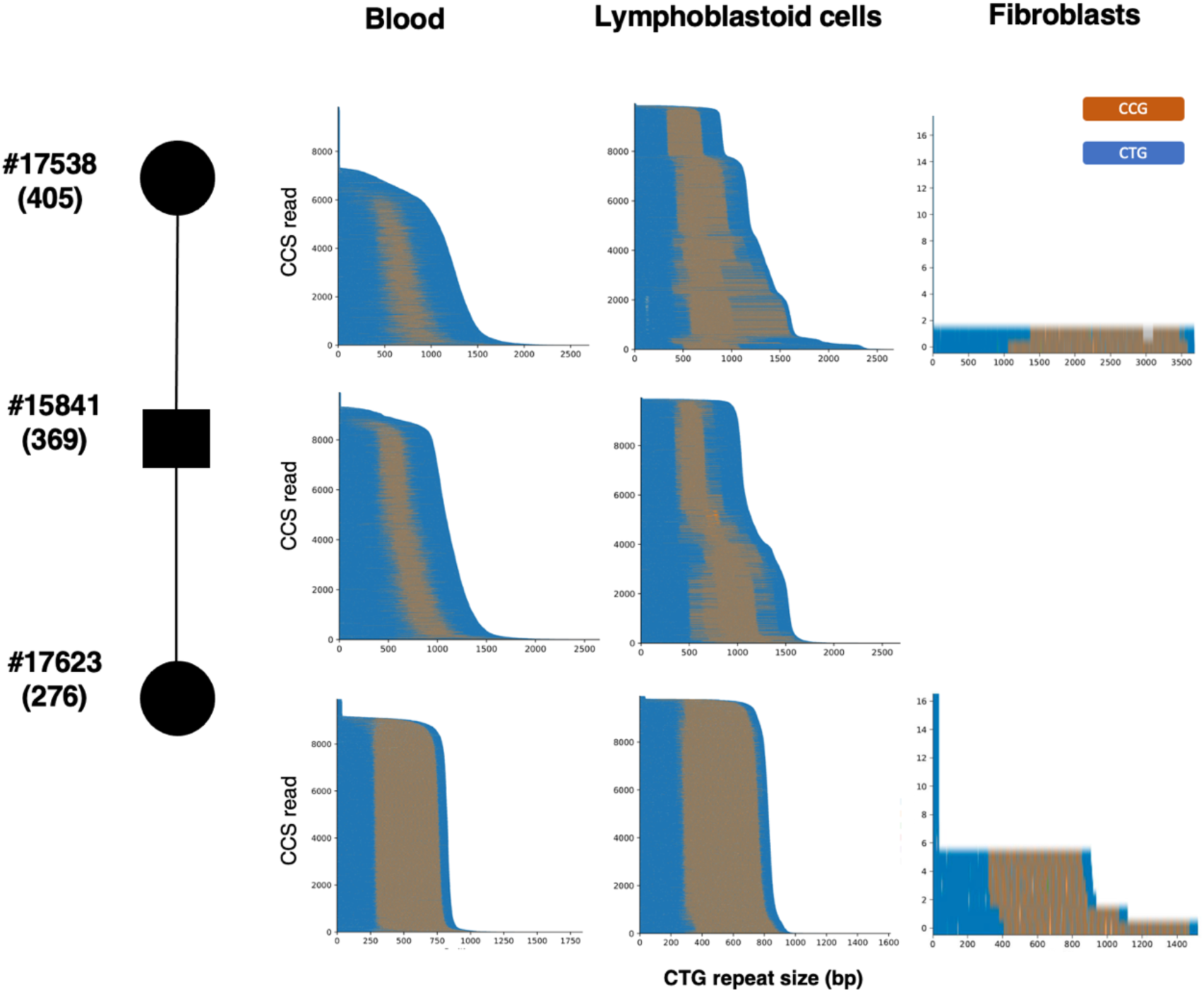
Waterfall plots highlighting the expansion structures at DM1 locus in blood, lymphoblastoid cell lines, and primary fibroblasts. For each plot, the x-axis shows the repeat expansion length in base pairs. The y-axis displays all the reads obtained after sequencing. CTG repeats are depicted in blue, and CCG repeats are shown in orange. The highest peaks correspond to the normal allele. On the left side of the figure, a portion of the pedigree from the atypical DM1 family analysis is displayed, following standardized human pedigree nomenclature (Bennet et al., 2008). The estimated size of the expanded allele in blood is shown in parentheses.

**Table 1:** Long-read sequencing metrics for individual samples #17538, #15841, and #17623. Coverage was determined by the number of high accuracy long reads.

| Samples | Methods | Coverage<br>(Expanded allele) | Size of the<br>expansion<br>(Median) | Interruptions<br>(Median) | % of<br>interruptions<br>(Median) | Age at<br>sample |
| --- | --- | --- | --- | --- | --- | --- |
| **Blood** |  |  |  |  |  |  |
| **17538** | Amplicon | 7280 | 405 (Max 899) | 52 CGG (Max 241) | 12,6 (Max 40) | 75 |
| **15841** | Amplicon | 9305 | 369 (Max 885) | 50 CCG (Max 215) | 13,7 (Max 39,8) | 58 |
| **17623** | Amplicon | 9167 | 276 (Max 613) | 79 CCG (Max 236) | 28,6 (Max 44) | 38 |
| **Lymphoblastoid**<br>**cell lines** |  |  |  |  |  |  |
| **17538** | Amplicon | 9863 | 401 (Max 890) | 85 CGG (Max 265) | 21,3 (Max 45,2) | 75 |
| **15841** | Amplicon | 9901 | 386 (Max 895) | 72 CCG (Max 219) | 18,5 (Max 46,5) | 58 |
| **17623** | Amplicon | 9809 | 276 (Max 537) | 81 CCG (Max 176) | 29 (Max 44,2) | 38 |
| **Fibroblasts** |  |  |  |  |  |  |
| **17538** | No-Amp-V1 | 2 | 1183 | 367 | 31 | ND |
| **17623** | No-Amp-V1 | 6 | 306 | 92 | 30 | ND |

Recently, we collected lymphoblast cell lines from each individual to analyze the dynamics of CCG interruptions in this cell type using HiFi long-read amplicon sequencing. Characterization of the DM1 locus revealed distinct cell populations in individuals #17538 and #15841, with differences in CCG interruption patterns and expansion sizes between them (Figure 1). Using non-amplification targeted long-read sequencing, we confirmed that the number and location of interruptions vary in lymphoblast cell line of #15841, indicating that the variation is not due to the amplification approach bias (Supplementary 1). In contrast, the CNG repeat expansion pattern in #17623 is consistent with the stable pattern observed in this individual’s blood, where the expansion remains highly stable. As observed in blood, 29% of CCG interruptions are observed in the DM1 expansion. To enhance our study, we analyzed the DM1 locus in primary fibroblasts derived from individuals #17538 and #17623. Characterization of the DM1 locus revealed that the number of interruptions varies in this cell type, with more significant variation noted in highly proliferative transformed cell lines. (Table 1 and Figure 1). The number and location of CCG interruptions appear to be variable not only within the same tissue but also between different cell types, suggesting that interruptions may vary across tissues in DM1 patients

### 2- The number of CCG repeats is not strictly correlated with the size of the repeat expansion

First, we examined the distribution of expansion sizes and CCG interruptions in the blood samples of individuals #17538, #15841, and #17623 (Figure 2A). The distribution of expansion size and CCG repeats are similar in all patients, with individual #17623 showing the most stable CTG and CCG somatic mosaicism. Therefore, we conducted an analysis on the relationship between the frequency of CCG interruptions and the magnitude of expansion in the blood samples of these individuals (Figure 2B). No significant correlation was found between the number of CCG interruptions and expansion size in individuals #17538 and #15841. However, individual #17623 showed a slightly positive correlation, with an R-squared value of 0.79, suggesting that the largest expansions are composed of a greater number of CCG triplets in the individual with the most stable expansion.

**Figure 2:**
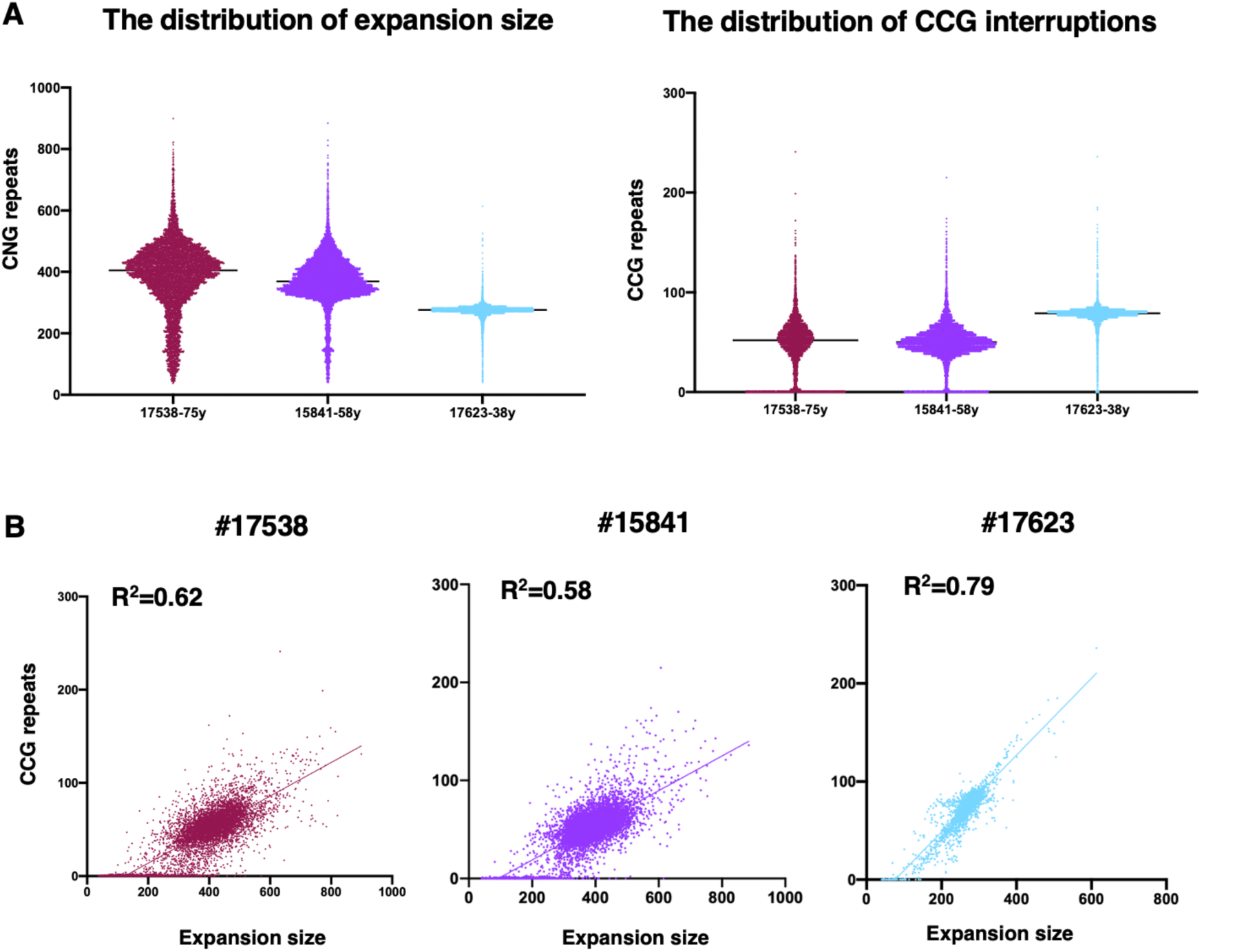
Dynamics of CNG repeat expansion and CCG interruptions in DM1 patients. **A:** On the left, the graph depicts the distribution of CNG repeat sizes for each HiFi read generated by the PacBio algorithm. The y-axis shows the number of CNG repeats, with each dot representing the size of CNG repeats in a single HiFi read. On the right, the graph illustrates the distribution of CCG interruptions for each HiFi read. The y-axis represents the number of CCG repeats, with each dot corresponding to the number of CCG repeats in a single HiFi read. The black bar indicates the median. A total of 7,000–9,200 single-molecule HiFi reads of the expanded allele were analyzed. **B:** The graph shows the correlation between the number of CCG repeats and the size of the expansion. A simple linear regression analysis was performed to determine the R-squared value, which is displayed in the graph. For each plot, the x-axis represents the size of the expansion in triplet repeats, and the y-axis displays the number of CCG repeats.

### CCG interruptions are methylated in DM1 patients

To explore DNA methylation (DNAme) at the DM1 locus, we conducted long-read free amplification sequencing (PacBio’s PureTarget) on the lymphoblastoid cell line of individual #15841, the only sample from the atypical DM1 family with sufficient DNA available for the study (Figure 3). Our findings revealed that CCG interruptions are methylated in this cell line. To further our investigation, we analyzed DNAme in the blood of members of a DM1 family with over 85% pure CCG repeats in the DM1 expansion, previously described in Tsai et *al.* (37). Specifically, we analyzed individual #N17, as described in Tsai et al., and his son, #C17, whom we recruited post-publication using whole genome sequencing (Table 2) (37). The results indicated methylated CCG repeats in both #N17 and #C17 (Figure 4A). However, DNA methylation exhibited variability in patient #C17 (prenatal DNA extraction). Notably, we have demonstrated for the first time that CCG-interrupted expanded alleles are methylated in analyzed DM1 patients, irrespective of the number and location of the repeats.

**Figure 3:**
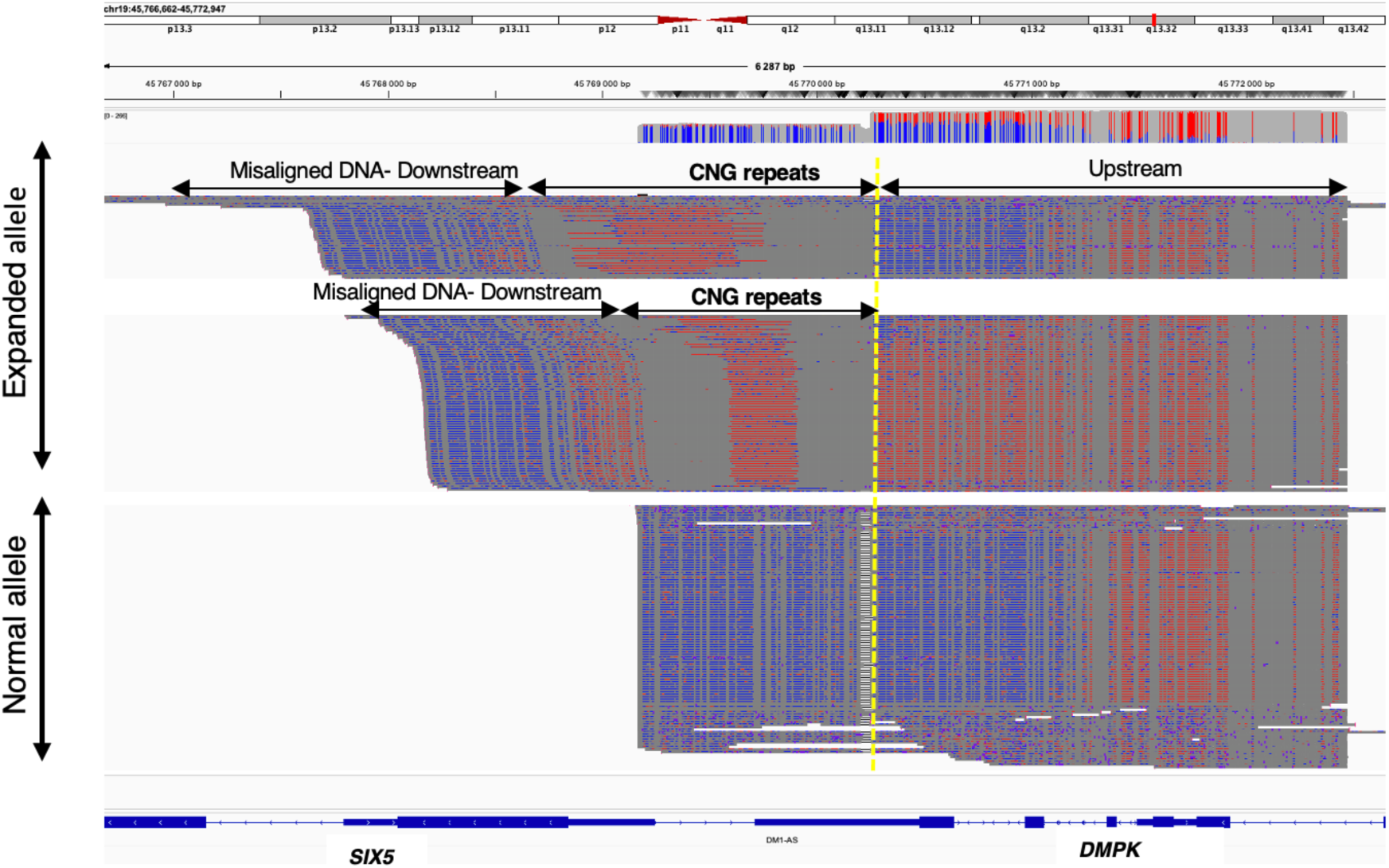
5-methylcytosine (5mC) methylation levels at the *DMPK* locus in #15841. IGV screenshot of the *DMPK* region in lymphoblastoid cell lines from sample #15841, showing the methylation status of each read. The normal allele (bottom) and the expanded allele (top) are separated by a black dotted line. The yellow dotted line delineates the downstream region from the 5’ end of the CNG repeat expansion. Each line represents an individual read. Unmethylated sites are colored blue, while methylated sites are colored red.

**Figure 4:**
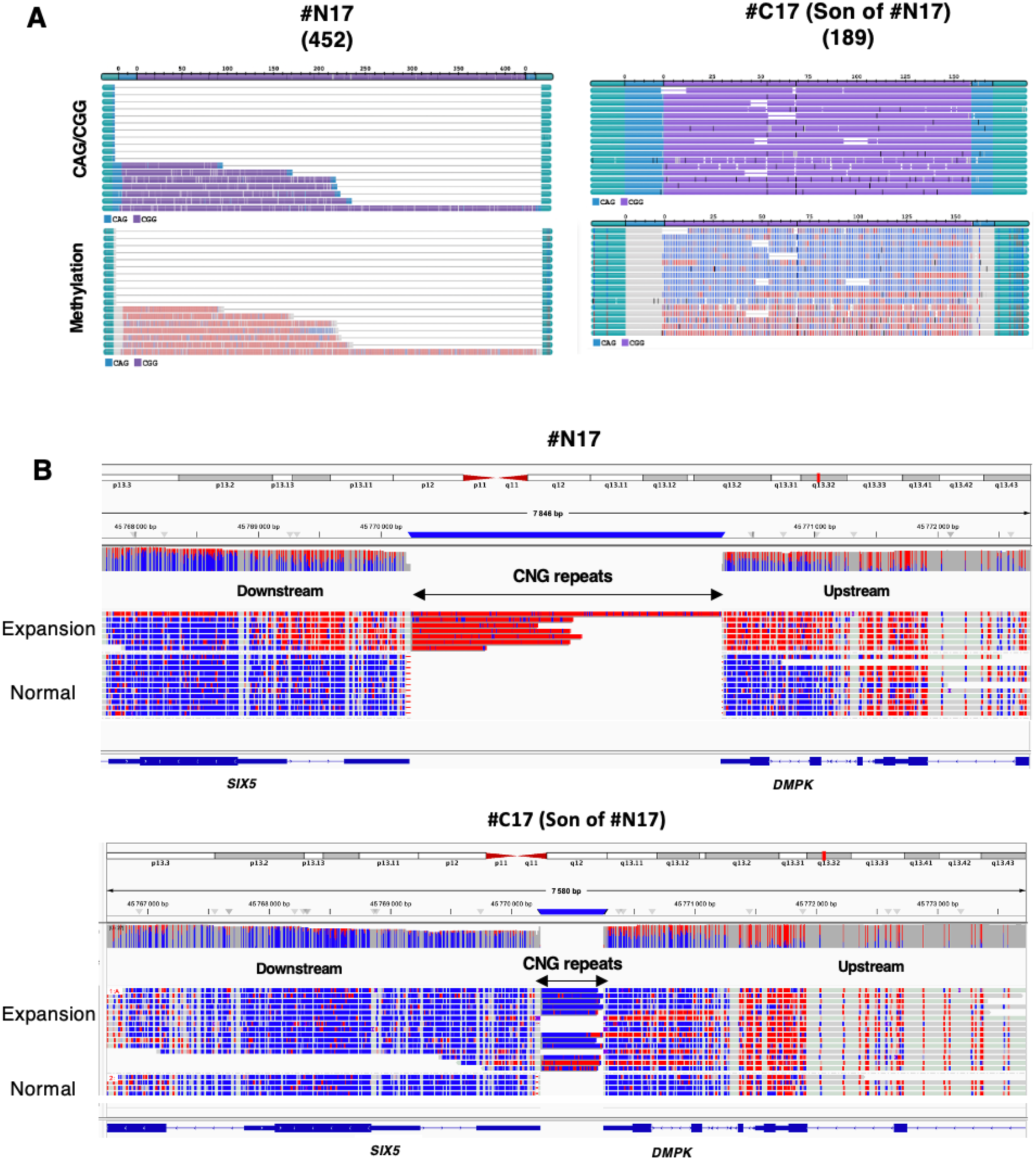
5-methylcytosine (5mC) methylation levels at the *DMPK* locus in samples #N17 and #C17. **A:** TRVZ plots of the CNG repeat for samples #N17 and #C17. The top panel shows the triplet repeat expansion structure, with CTG repeats represented in blue and CCG repeats in purple. The bottom panel displays CpG methylation, with unmethylated sites colored blue and methylated sites colored red. These plots illustrate the repeat alleles and reads aligning to them, along with optional methylation information. **B:** IGV screenshot of the *DMPK* region showing the methylation status of each read. The normal allele (bottom) and the expanded allele (top) are separated by a black dotted line. Each line represents an individual read, with unmethylated sites colored blue and methylated sites colored red. The estimated size of the expanded allele in blood is shown in parentheses.

**Table 2:**
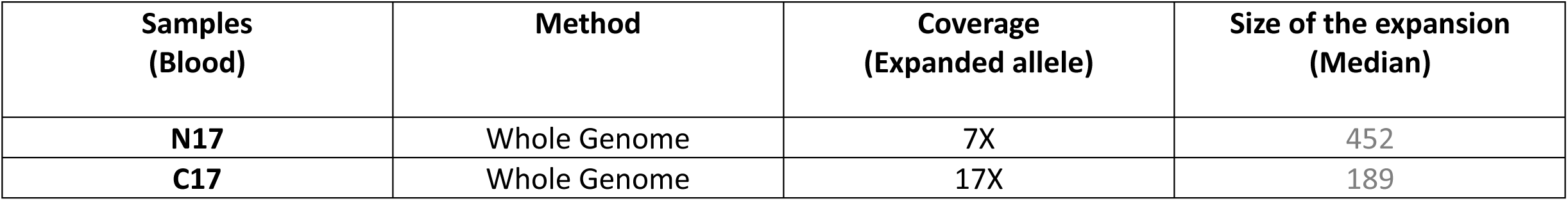
Long-read sequencing metrics for individual samples #N17 and #C17. Coverage was determined by the number of high accuracy long reads.

### 3- CCG interruptions are associated with hypermethylation within the 3’UTR of *DMPK*

To assess the impact of CCG interruptions on DNA methylation in the flanking regions of the *DMPK* locus in DM1 patients, we analyzed 5-methylcytosine (5mC) methylation pattern in the lymphoblastoid cell line (LCL) of individual #15841, as well as in samples from #N17 and #C17 (Figure 3 and 4B). In the #15841 LCL, two distinct populations were observed, showing varying DNA methylation patterns both upstream and downstream of the repeats (Figure 3). The highest methylation levels were found in the shortest CNG repeat expansion alleles, where both upstream and downstream regions were methylated. In #N17, aged 21 at the time of sampling, CCG interruptions induced methylation both upstream and downstream of the repeats. However, in the fetal sample #C17, the levels of DNA methylation in the flanking region vary, with the highest methylation levels observed upstream of the repeats (Figure 4B).

## Discussion

Sequence interruptions were first described in fragile X syndrome and have been identified in various trinucleotide repeat disorders (4,43). In the context of DM1, CCG, GGC, and CTC interruptions were first reported at the 3ʹ end of the CTG repeat expansion in DM1 patients (19,20). More recently, multiple CCG interruptions, as well as a single CAG repeat interruption at the 5ʹ end of the CTG repeat sequence have been observed in DM1 families (21,23). In contrast to the pattern exhibited by other trinucleotide repeats, the nature and number of CNG interruptions in the DM1 population display marked variability. The use of targeted long-read sequencing offers a unique opportunity to accurately characterize DM1 triplet repeat expansions and to better understand triplet repeat interruption behavior in DM1 patients. Using LRS innovative approach, we have demonstrated that the number of CCG interruptions can vary across generations and between cell types, despite a global stabilization of the expansion in DM1 patients. This finding complicates the understanding of the mechanisms involved in the atypical instability observed in patients carrying CCG-interrupted expansion alleles. Multiple studies have shown that interruptions like AGG, CAT, GGA, and CCG can affect DNA structure, methylation, and repeat instability in various diseases by altering repair and replication processes (29–34). All of these processes could be differently disrupted by the type, number, and location of interruptions. In this study, we focused on CCG interruptions in certain DM1 patients, analyzing 5mC methylation pattern both in the flanking regions of the DM1 repeats and within the repeats themselves. Notably, we have shown for the first time that CCG repeats are methylated in these patients, and this methylation was consistently associated with hypermethylation in the flanking sequences in all analyzed DM1 patients. For the upstream part, the results are more complex and depend on the model used. Our study highlights the intricate behavior of CCG interruptions in DM1, revealing significant variation across different tissues and even within the same tissue. The impact of CCG interruptions on DNA methylation in DM1 patients depends on multiple factors, including age, expansion size, tissue type, and the number and location of interruptions. This complexity is supported by previous studies comparing patients with interrupted versus pure CTG repeat tracts, which show varied CpG methylation patterns. For instance, Santoro et *al.* noted an altered methylation profile in patients with 400–900 interrupted CTG repeats, with increased downstream but no upstream CpG methylation whereas other studies have shown no change in DNA methylation profile within 3’UTR of *DMPK* (17,18,23,29). In one previous study, it has shown that an increased DNA methylation downstream of the repeat is associated with reduced somatic mosaicism (17). This finding supports the hypothesis that downstream hypermethylation in DM1 patients with CCG interruptions could explain the stabilization of CTG repeats observed in these individuals. Légaré et al. found that DNA methylation at two CpG sites downstream of the CTG repeat expansion is associated with respiratory failure and muscle weakness, regardless of CTG repeat length (17). Similarly, a 9-year longitudinal study of 115 DM1 patients indicated that DNA methylation levels downstream of the CTG repeat were linked to changes in cognitive features (44). Together, these studies underscore the importance of epigenetic modifications at the *DMPK* locus in the clinical manifestations of DM1.

Our study clearly highlights the complexity of DM1 genetics and the significant variability observed within the small cohort of DM1 patients we analyzed. These observations underscore the need for a more refined molecular approach in a large cohort of DM1 patients to better understand the mechanisms of repeat instability and its clinical manifestations. The next step involves large-scale analyses using advanced long-read sequencing technologies to improve genotype-phenotype correlations and enhance genetic counseling and prognosis for DM1 patients.

## Ethics approval

Participants gave informed consent to participate in the study before taking part.

## Acknowledgements

The authors would like to thank the DM1 patients and colleagues at the Myology Centre for Research, Curie Institute and PacBio for helpful discussions and comments. We specially thank Maeva Froidevaux, Eliane Gardais, Sana Ghazran, Safaa Saker-Delye and Nadaj Pakleza Aleksandra. This work was supported by the Institut National de la Santé et de la Recherche Médicale, Sorbonne Université and Institut de Myologie.

---

**Supplementary 1:**
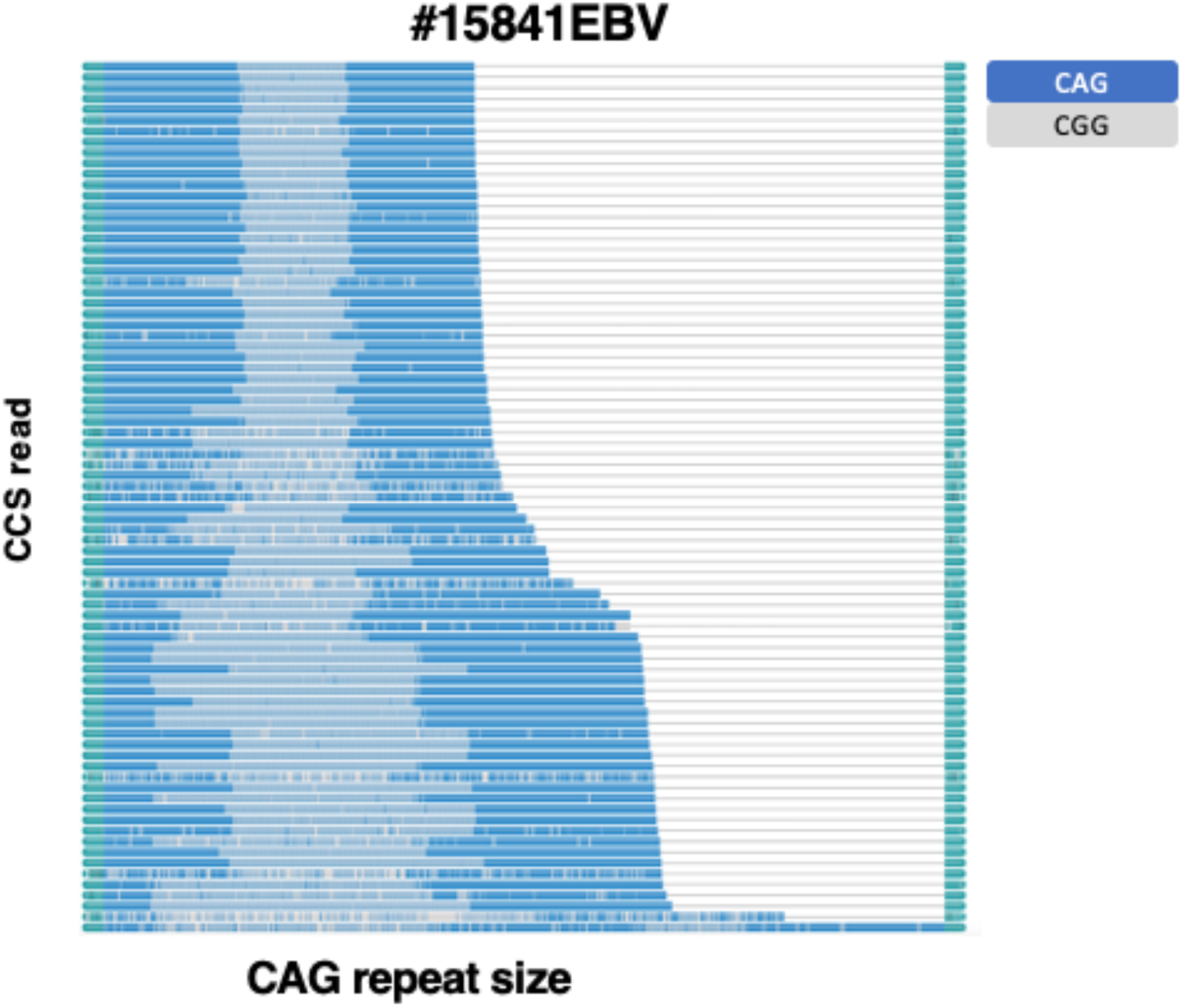
TRVZ plot of the CNG repeat for #15841 (Lymphoblastoid cells). CTG is represented in blue and CCG repeat in gray.

